# Climate, habitat, and geographic range overlap drive plumage evolution

**DOI:** 10.1101/375261

**Authors:** Eliot T. Miller, Gavin M. Leighton, Benjamin G. Freeman, Alexander C. Lees, Russell A. Ligon

**Affiliations:** Cornell Lab of Ornithology, 159 Sapsucker Woods Rd., Ithaca, NY 14850, USA; Department of Zoology, University of British Columbia, #4200-6270 University Blvd, Vancouver, BC V6T1Z4, Canada; School of Science & the Environment, Manchester Metropolitan University, Manchester, UK

## Abstract

Organismal appearances are shaped by selection from both abiotic and biotic drivers ^1–5^. For example, Gloger’s rule describes the pervasive pattern that more pigmented populations are found in more humid areas ^1,6,7^, and substrate matching as a form of camouflage to reduce predation is widespread across the tree of life ^8–10^. Sexual selection is a potent driver of plumage elaboration ^5,11^, and species may also converge on nearly identical colours and patterns in sympatry, often to avoid predation by mimicking noxious species ^3,4^ To date, no study has taken an integrative approach to understand how these factors determine the evolution of colour and pattern across a large clade of organisms. Here we show that both habitat and climate profoundly shape avian plumage. However, we also find a strong signal that many species exhibit remarkable convergence not explained by these factors nor by shared ancestry. Instead, this convergence is associated with geographic overlap between species, suggesting strong, albeit occasional, selection for interspecific mimicry. Consequently, both abiotic and biotic factors, including interspecific interactions, are potent drivers of phenotypic evolution.

---

Birds are frequently studied for the insights they can offer to our understanding of phenotypic evolution ^11,12^ Yet, we still do not understand how factors such as climate, substrate matching, and sexual selection interact to influence phenotypic evolution across a large clade of birds (or any other large organismal radiation, for that matter). Here, we employ a novel phylogenetic comparative framework, coupled with remote-sensing data and a large citizen science dataset, to examine the combined effects of climate, habitat, evolutionary history, and social selection for mimicry on plumage pattern and colour evolution in woodpeckers. This diverse avian clade of 230 species is an excellent group to examine how external phenotypes evolve because they occupy a broad range of climates across many habitats globally, and display patterns of rapid divergence in appearance, as well as striking convergences. A complete, time-calibrated phylogeny, which we incorporated into our analyses here, recently highlighted the magnitude of some of these convergences ^13^. Woodpeckers have evolved a wide range of plumages; from species with boldly marked black and white patterns and others with large bright red patches, to other species that are entirely dull olive (Fig. 1). This variation provides the raw material with which to disentangle the contribution of the various abiotic and biotic factors that may drive plumage evolution. Moreover, there are several cases of ostensible plumage mimicry within woodpeckers ^14,15^; whether such cases represent biotic or socially driven convergence, or whether they can be explained by shared ancestry, climate, or habitat space remains untested.

**Fig. 1.**
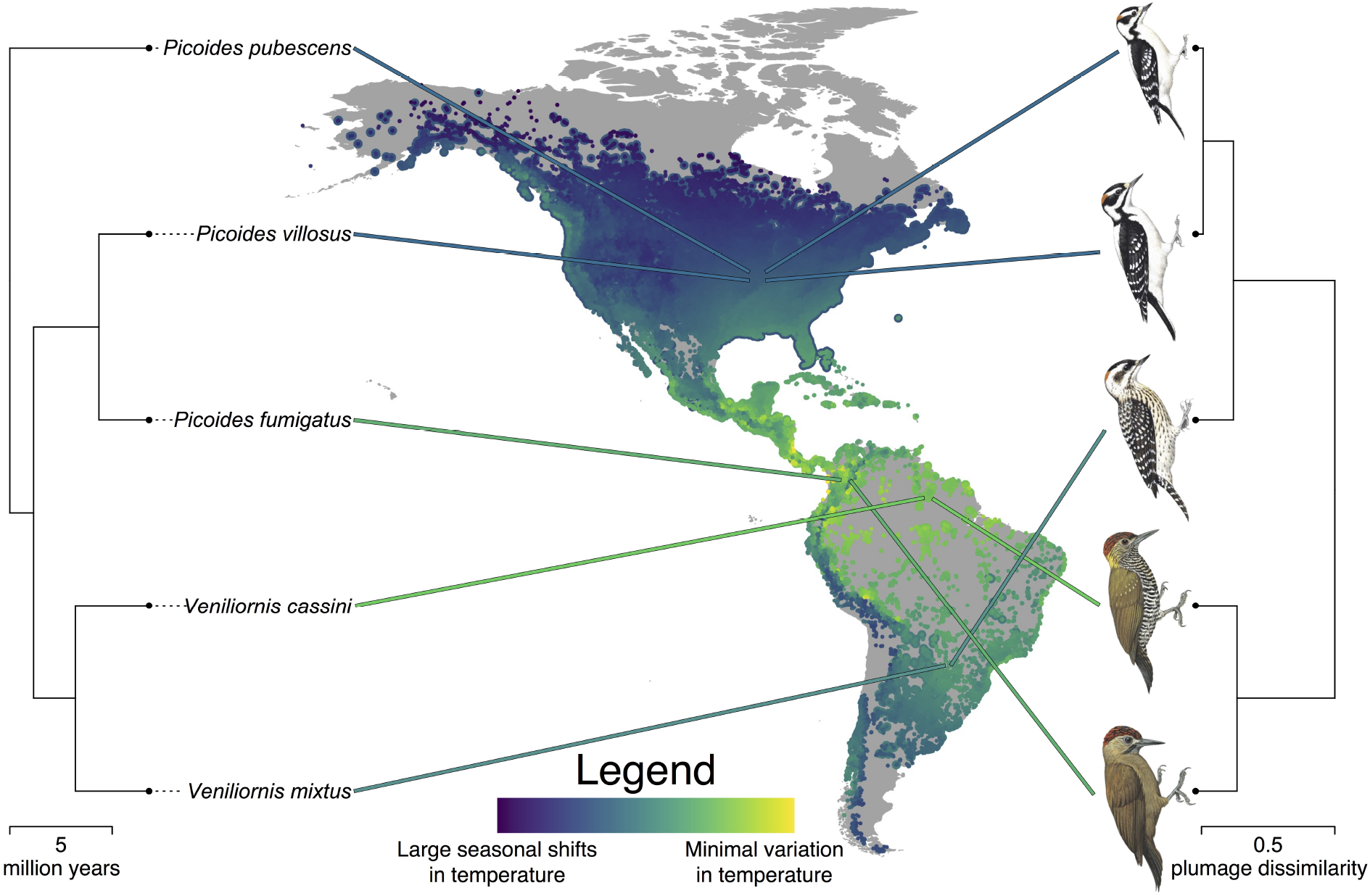
Evolutionary relationships and plumage similarity among exemplar species that shared a common ancestor ~6.5 mya. Climate partially determines variation in woodpecker plumage. Lines lead from tips of phylogeny (left) to centroid of each species’. geographic distribution, and are coloured according to mean climate regime of each species. The colour scale depicts a gradient from warm (yellow) to seasonally cold regions (blue). eBird records for these species are plotted in the same colours as large points on the map. All other eBird woodpecker records are overlaid as smaller points and coloured similarly. Plumage dendrogram (right) shows the plumage dissimilarity relationships among the same set of species. *Veniliornis mixtus*, long classified as a member of *Picoides*, is inferred to have invaded seasonal climates in the southern hemisphere, and accordingly evolved bold black and white plumage. *Picoides fumigatus*, long classified as a member of *Veniliornis*, is inferred to have invaded warm climates near the equator, and accordingly evolved dark, subtly marked plumage. *Picoides pubescens* and *P. villosus* are rather distantly related but largely sympatric; they are inferred to have converged on one another in plumage above and beyond what would be expected based on shared climate, habitat, and evolutionary history. Illustrations © HBW Alive/Lynx Edicions.

We used multidimensional-colour and pattern-quantification tools to measure species’ colouration and patterns, quantifying species’ plumages from a standardized source (Figs. 2–3) ^11,16,17^ Evidence suggests that pattern and colour are likely processed separately in vertebrate brains, with achromatic (i.e., luminance) channels being used to process pattern information ^18^ and differential stimulation of cones encoding chromatic information ^19^. While both plumage colour and pattern are inherently multivariate, we reduced this complexity into a composite matrix of pairwise species differences. We could then address whether purported convergences were a mere by-product of shared evolutionary history or, if not, which of the aforementioned factors were responsible. We incorporated the potential for interactions between pairs of species into the analysis by quantifying pairwise geographic range overlap using globally crowd-sourced citizen science data from eBird ^20^; species in complete allopatry have no chance of interacting, while increasing degrees of sympatry should correlate with the probability of evolutionarily meaningful interactions.

**Fig. 2.**
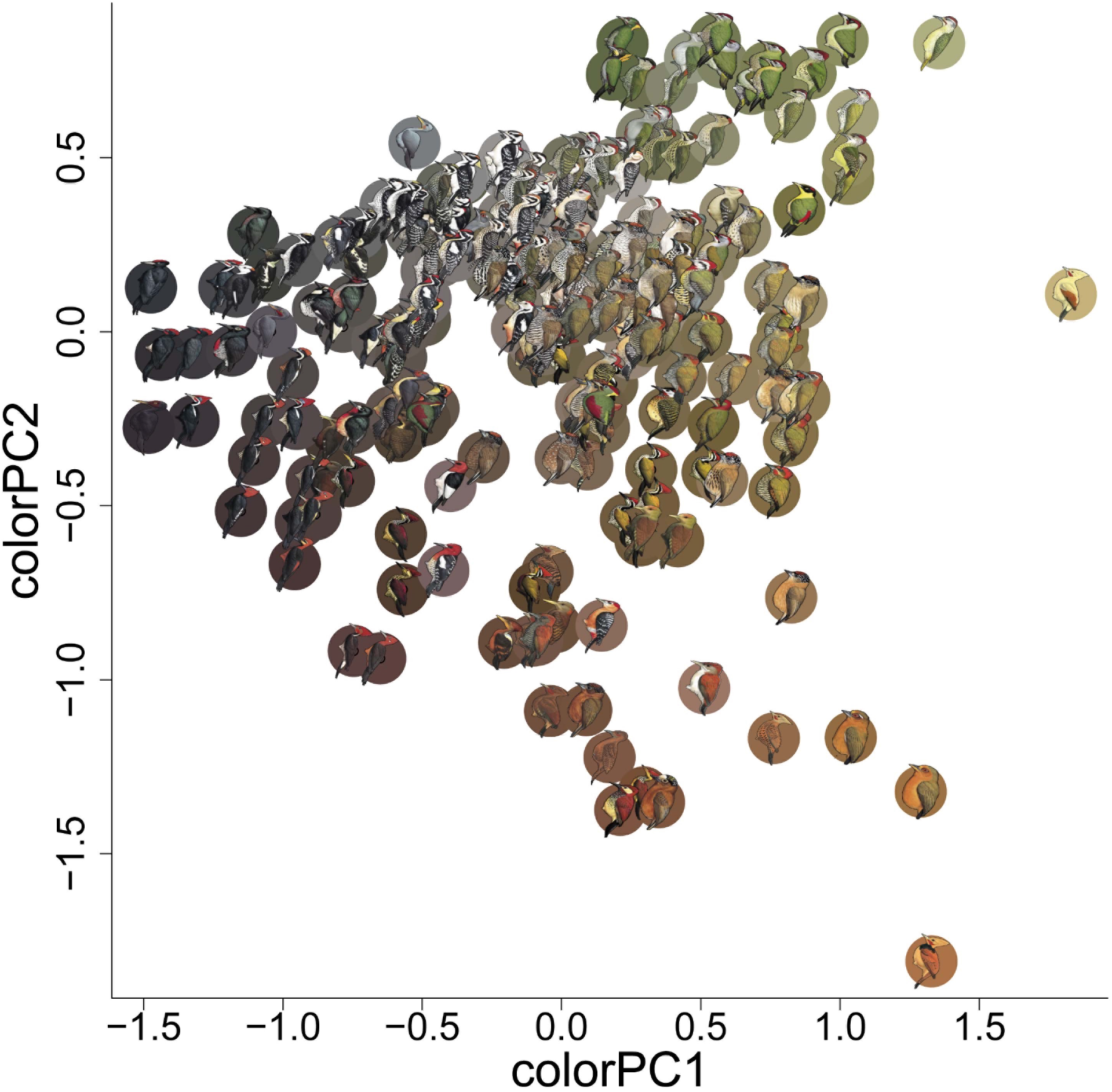
Principal components analysis (PCA) of species-averaged woodpecker colour values, used for species-level analyses (e.g., Fig. 4.). Principal component one (colourPC1) explains 45% of the variation in measured colour scores. Higher PC1 scores correspond to greater luminance values, and more yellow and less blue. Principal component two (colourPC2) explains an additional 36% of variation in overall colour scores. Higher PC2 scores correspond to more green and less red colouration. Coloured circles behind each woodpecker species correspond to the average CIE L*a*b scores for the 1,000 randomly drawn colour samples from that species. Illustrations © HBW Alive/Lynx Edicions.

**Fig. 3.**
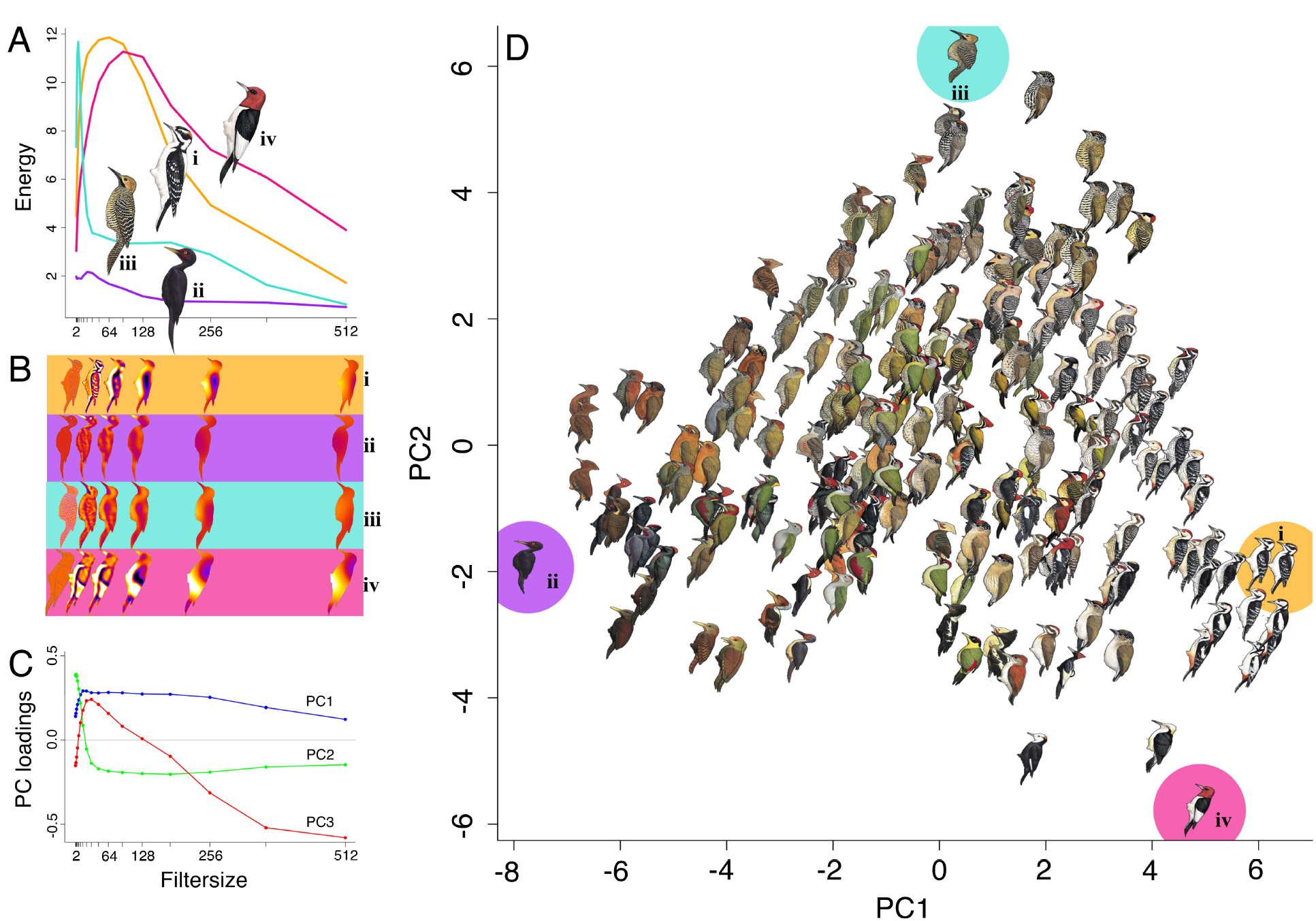
Major axes of plumage pattern variation quantified using granularity analysis, then summarized with a principal components analysis (PCA) for species-level approaches. (a) Pattern energy spectra for exemplar species characterized by maximally divergent PC scores (i = *Picoides villosus*, ii = *Mulleripicus funebris*, iii = *Colaptes fernandinae*, iv = *Melanerpes erythrocephalus*). These show information on relative contributions of different granularities to overall appearance. (b) Pattern maps of exemplar species enable visualization of energy at subsets of isotropic band-pass filter sizes (2, 32, 64, 128, 256, and 512 pixels). (c) Principal component loadings of pattern energy spectra across filter size for all species reveals how pattern elements of different sizes influence PC scores. (d) Pattern PC1 and PC2, collectively, account for 82.1% of variation across woodpeckers. Exemplar species (i-iv) illustrate extreme variation in PC1: (i) exhibits high energy scores across most pattern element sizes, with small, medium, and large pattern elements; (ii) has low energy scores across the spectrum, with few pattern elements of any size. And, extreme variation in PC2: (iii) has many small pattern elements and few of any other sizes; (iv) has only medium and large size pattern elements. Illustrations © HBW Alive/Lynx Edicions.

Comparing species-level pairwise distance matrices revealed that variation in climate (*r* = 0.055, *p* = 0.006), habitat (*r*
 = 0.106, *p* = 0.007) and, to a lesser degree, phylogenetic relationships (*r* = 0.001, *p* = 0.015) are all correlated with woodpecker plumage similarity scores. Overall, woodpecker species in similar climates and habitats tend to look alike, even after accounting for shared ancestry. However, we also found that close sympatry was a strong predictor of plumage similarity for the most similar-looking species pairs (Extended Data Fig. 1). We interpret this result as evidence for widespread plumage mimicry *per se,* separate from any broader patterns of environmentally-driven plumage convergence. Following this result, we developed a novel method to identify the species pairs that powered this striking result. Many previously suggested mimicry complexes were, in fact, responsible, including the Downy-Hairy Woodpecker system (Fig. 1) ^21^, repeated convergences between members of *Veniliornis* and *Piculus* ^22^, *Dinopium* and *Chrysocolaptes, Dryocopus* and *Campephilus* ^23^, and the remarkable convergence of *Celeus galeatus* on the latter two genera ^15^. Collectively, this combination of distance matrix-based analyses provides powerful understanding of the importance of various factors in driving evolutionary patterns of mimicry and divergence. To gain further insight into the evolutionary drivers of particular colours and patterns, we delved deeper with species-level phylogenetic comparative approaches.

We found that precipitation consistently drives global patterns of plumage pigmentation and patterning in woodpeckers. We show that darker species tend to inhabit areas of higher precipitation (phylogenetic generalized least squares [PGLS], *r*^2^ = 0.08, *p* < 0.001), supporting Gloger’s rule ^1^. While not unexpected, this is the first time that Gloger’s rule has been quantitatively assessed for a large, globally distributed clade. The mechanism underlying Gloger’s rule remains debated, but may include a response to increased predation pressure in the tropics ^24^ via background matching ^25^, or defence against feather-degrading parasites ^26^. In addition to providing support for Gloger’s rule, wherein precipitation is correlated with dark plumage, we show that precipitation is also associated with reduced patterning (PGLS *r*^2^ = 0.15, *p* < 0.001). These results could be interpreted as support for either or both proposed mechanisms. For example, there are some boldly marked woodpecker species in humid areas, but they invariably achieve these conspicuous phenotypes with minimal use of white plumage, suggesting an evolutionary trade-off wherein Gloger’s rule is due, at least in part, to the ability of melanin to forestall feather wear ^26^. Alternatively, unconcealed large, white plumage patches might simply subject humid forest-dwelling birds to evolutionarily unacceptable levels of predation. While additional research is necessary to pin down the mechanism(s) responsible, our results expand the generality of Gloger’s rule and show that it may be involved in phenotypic convergence among disparate lineages inhabiting similar forests. For example, *Micropternus brachyurus* is not closely related to the members of *Celeus,* as was long thought, and instead simply occupies Asian forests of similar physiognomy to those inhabited by some Neotropical *Celeus.*

Seasonality, in addition to total precipitation, also exerts significant influence on woodpecker plumage. The gradient from dark-to light-plumaged woodpeckers (colorPC1) was best explained by a model that included body mass, latitude, and seasonality in precipitation. Darker birds are larger, are found at lower latitudes, and in climates that receive much precipitation throughout the year (PGLS *r*^2^ = 0.08, *p* < 0.001, Fig. 4a). The gradient from red to green plumaged woodpeckers (colorPC2) was best explained by a model that included variation in temperature seasonality, and included the dichotomy between open habitats and closed forests. Specifically, green birds tend to be found in climates that experience seasonal temperature fluctuations, and in open habitats (PGLS *r*^2^ = 0.07, *p* < 0.001, Fig. 4b). Moreover, boldly marked birds tend to be found in seasonal climates, open habitats, and temperate forests, and the gradient from boldly to subtly marked birds was best explained by a model that included variation in temperature seasonality, and the dichotomies between open habitats and closed forests, and between dense tropical forests and temperate forests (PGLS *r*^2^ = 0.15, *p* < 0.001, Fig. 4c). We had suspected that variation along the gradient from species with large plumage elements to those with barring and spotting (pattern PC2) might be associated with sexual selection, but after accounting for body mass, pattern PC2 was not associated with sexual size dimorphism; instead, smaller birds simply tended to be more finely marked (PGLS *r*^2^ = 0.04, *p* = 0.01, Fig. 4d). This plumage pattern gradient was not closely associated with any climate or habitat gradient, and we could find no evidence that sexual selection shapes woodpecker plumage in any consistent way across taxa.

**Fig. 4.**
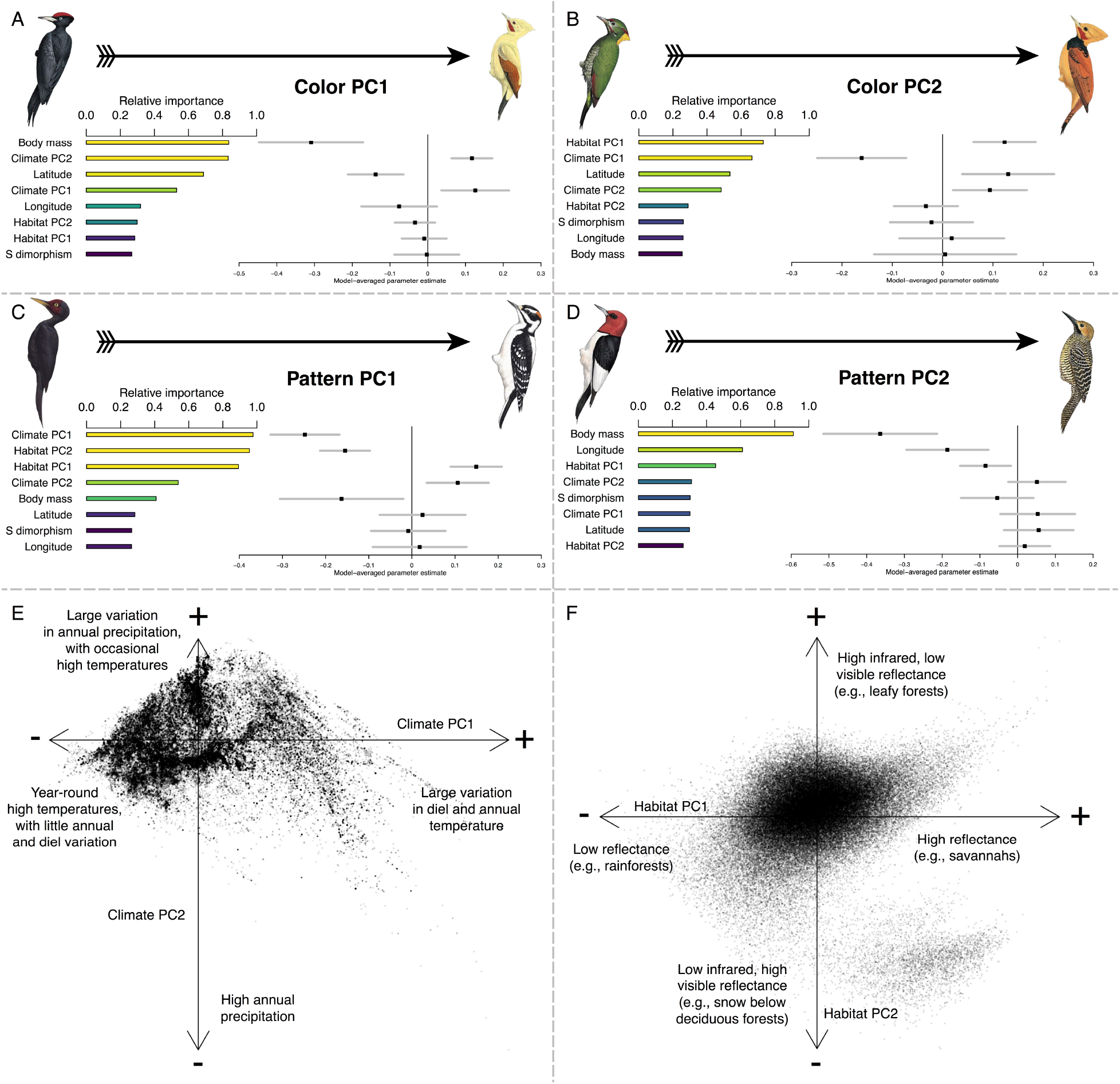
Variable importance scores and model-averaged parameter estimates from phylogenetic generalized least squares regressions. These quantify how colour and pattern vary as a function of climate, habitat, body mass, sexual size dimorphism, latitude and longitude, with summaries of the climate and habitat principal component analyses (PCA). Model-averaged *p*-values of explanatory factors are colour-coded from yellow to blue; only factors with *p*-values < 0.05 are coloured yellow and discussed here. (a) Dark birds are heavier and occur in wetter climates. (b) Greenish (as opposed to reddish) birds are found in more open habitats. (c) Less-patterned birds are found in aseasonal climates, open habitats, and temperate forests. (d) Birds patterned in large plumage elements, such as large colour patches, tend to be larger in body size. (e) Climate PCA results, illustrating the distribution of woodpeckers in climate space, with qualitative descriptions of the first two PC axes. (f) Habitat PCA results, showing the distribution of woodpeckers across global habitats, with qualitative descriptions of the first two PC axes. Illustrations © HBW Alive/Lynx Edicions.

Although climate and habitat appear responsible for some of the convergence in external appearance in woodpeckers, our analyses confirmed the decades-old suggestions ^27^ that some species have converged above and beyond what would be expected based only on selection pressures from the environments they inhabit. Sympatry, a proxy for the likelihood of evolutionarily meaningful interspecific interactions, was a strong predictor of plumage similarity for species exhibiting large geographic range overlaps (Extended Data Fig. 1). Given that sympatry itself is implicated in plumage convergence, the evidence indicates that such convergence is true mimicry, i.e., phenotypic evolution by one or both parties in response to a shared signal receiver ^3,4^. Indeed, our study almost certainly underestimates the degree to which close sympatry leads to mimicry in woodpeckers: some mimetic dyads are well known to track one another at the subspecific level. Recent taxonomic revision of *Chrysocolaptes* ^28^ that is not yet matched by equivalent efforts in *Dinopium* means that additional parallel mimicry in this complex was not entirely accounted for in our study (e.g., the extraordinary convergence between the Sri Lankan endemics C. *stricklandii* and *D. benghalense psarodes).* There are two contested questions regarding plumage mimicry: whether it truly occurs ^23,27,29^ and, if it does, what process(es) drive the pattern ^30–33^. Here we have shown that plumage mimicry does indeed occur and is pervasive across the woodpecker evolutionary tree, indicating that the processes deserve further study.

Few studies ^21,34^ have empirically demonstrated that convergence on the scale which we document among woodpeckers is not exclusively a product of shared evolutionary history. Nevertheless, several mechanisms have been proposed to explain convergent evolutionary trajectories of external appearance ^27,31,32,35,36^. Recently, it has been shown ^33^ that the smaller species in plumage mimicry complexes may derive a benefit by fooling third parties into believing they are the socially dominant model species, and currently this remains the only empirically supported hypothesis. An open question is how distantly related lineages genomically achieve plumage convergence. It is unclear whether multiple mutations are required, each of which increases the degree of plumage convergence, or if selection acts on genetic modules controlled by fewer loci that shape overall plumage and are shared across all woodpeckers, or if possibly rare hybridization events between sympatric species have resulted in adaptive introgression of relevant plumage control loci ^22,37^.

In summary, we provide the first demonstration of selection promoting the evolution of mimicry at a broad phylogenetic scale. Additionally, we provide novel, comprehensive tests of longstanding hypotheses about the relative importance of abiotic and biotic factors to the evolution of organisms’ external appearances. Because we incorporated enormous variation in plumage, climate, habitat, and geographic range overlap into our phylogenetic comparative framework, our explanatory power was limited. Nevertheless, our simplified univariate PGLS approach corroborated the finding that climate and habitat can drive phenotypic evolution in predictable ways across a globally distributed clade of birds. Moreover, we showed that the convergence predicted by shared climate, habitat, and evolutionary history is insufficient to explain the prominent, geographically-widespread convergent events that we detected between closely sympatric, but distantly related woodpecker species. The modern union of comprehensive, time-calibrated molecular phylogenies, massive distributional databases like eBird, and powerful computing techniques like pattern analysis enables rigorous testing of hypotheses first put forth centuries ago. Within this framework, our results provide quantitative details of the abiotic and biotic factors, including interspecific signalling, that act as potent drivers of phenotypic evolution.

## METHODS

### Taxonomic reconciliation and creation of a complete woodpecker phylogeny

A time-dated phylogeny containing nearly all known woodpecker species was recently published by Shakya and colleagues ^13^. As described below, we used illustrations from the Handbook of the Birds of the World Alive ^16^ to quantify woodpecker plumage, and we used eBird ^20^, a massively crowd-sourced database, to define spatial, climate, and habitat overlap between species. Each of these references uses a slightly different taxonomy. Our goal was to use the species-level concepts from the most recent eBird/Clements taxonomy ^38^ as our final classification system.

To reconcile these three taxonomies (HBW, Shakya et al., and eBird/Clements), we first identified any taxon that was represented by multiple tips in the Shakya et al. (2017) tree. All of the species represented by multiple tips were monophyletic except one: D. *moluccensis.* However, eBird only recognizes a single taxon of this species. Thus, we simply dropped all but one instance of all species represented by multiple tips in the Shakya tree. In the process, we noted that *Dendrocopos medius* is represented by multiple tips in the Shakya tree, but the second instance is listed under *Leiopicus medius*. The two sequenced specimens are monophyletic, so we dropped the latter. We then identified all named woodpecker taxa in eBird. We allocated these to their respective terminals in the Shakya tree manually, to ensure accurate matches. For example, *Picus miniaceus* eBird records were allocated to *Chrysophlegma miniaceum* from the Shakya tree, and the tip renamed accordingly. This process made it clear which species, as recognized by eBird, were missing from the Shakya tree.

Twenty-one such missing taxa were identified: *Picumnus fuscus, P. limae, P. fulvescens, P*. *granadensis*, *P. cinnamomeus, Dinopium everetti*, *Gecinulus viridis*, *Mulleripicus fulvus*, *Piculus simplex*, *Dryocopus hodgei*, *Melanerpes pulcher*, *Xiphidiopicus percussus*, *Veniliornis maculifrons*, *Dendrocopos analis*, *Dendrocopos ramsayi*, *Colaptes fernandinae*, *Chrysocolaptes festivus*, *C. xanthocephalus*, *C. strictus*, *C. guttacristatus*, and *C. stricklandi*. We added these using the R package *addTaxa* ^39^, and taxonomic hypotheses outlined in previous work (reviewed in ^13^). Eighteen of these taxa have fairly precise hypothesized taxonomic positions which we were able to leverage to carefully circumscribe where they were bound into the tree. As an example, *Dinopium everetti* was recently split from *D*. *javanense*, so it was simply added as sister to the latter species in the final tree. The precise phylogenetic positions of the remaining three taxa are less well known. For these, we first added *C. fernandinae* as sister to *Colaptes sensu stricto* (as previously found ^40^), then added *Piculus simplex* into the clade *Piculus* C *Colaptes,* as previous work showed some members of the former genus to actually belong to the latter ^40^. We added *X. percussus* as sister to *Melanerpes striatus* (reasons summarized in Shakya et al. (2017); and we added P. *cinnamomeus* into *Picumnus* while ensuring that the Old World P. *innominatus* remained sister to the rest of the genus (it is very likely the New World *Picumnus* form a clade). Finally, we dropped a few tips from the phylogeny that represent splits not currently recognized by eBird: *Celeus ochraceus, Melanerpes santacruzi,* and *Picus sharpei.* The resulting tree contained 230 species. For each of these taxa, we identified the illustration that best represented it in the Handbook of the Birds of the World Alive. When the latter recognized multiple subspecies for a given taxon from the final tree, we used the nominate subspecies as our unit of analysis for colour and pattern (see below).

### Quantifying plumage colour and pattern from illustrations

We calculated plumage colour and pattern scores for males of 230 species of woodpeckers using digital images of colour plates obtained from *The Handbook of the Birds of the World Alive* ^16^. Each image was imported to Adobe Photoshop (Adobe Inc. San Jose, CA) at 300 dots per inch, scaled to a uniform size, and saved as a Tagged Image File (.TIF). Following creation of .TIF files, we ran a custom macro in ImageJ ^41^ to sample the red (R), green (G), and blue (B) pixel values for each of 1,000 random, 9-pixel-diameter circles from each woodpecker image (macro code will be made available with data upon acceptance). RGB values were transformed to CIELAB coordinates, which is an approximately perceptually-uniform colour space (distance between points is perceptually equivalent in all directions) ^42,43^. To calculate pairwise colour dissimilarity scores, we plotted the 1,000 colour measurements from the first species (e.g., species A) in 3-dimensional CIELAB space, as well as the 1,000 measurements for the second species (e.g., species B) in the dyadic comparison. We then calculated the average Mahalanobis distance ^44^ between the colours representing species A and the colours representing species B. We repeated this process for every possible combination (26,335 unique dyadic combinations) to generate an overall colour dissimilarity matrix. Additionally, to facilitate a more in-depth investigation of the underlying variation in colour among species, as well as how such variation is related to environmental, genetic, and geographic influences we conducted principal components analysis (PCA) on all 230,000 colour measurements. Following PCA, we averaged principal component (PC) scores for each species to create mean PC scores describing the average colour values for each species (Fig. 2). PC1 describes a dark-to-bright continuum, as well as a blue-to-yellow continuum (high loadings for L* and b*; Extended Data Table 1; Fig. 2), while PC2 primarily describes a red-to-green continuum (high loadings for a*; Extended Data Table 1; Fig. 2).

We conducted pattern analyses on the same, scaled .TIF files for each species in ImageJ 41. First, we split each image into R, G, B slices and then used the G layer for pattern analysis because this channel corresponds most closely to known avian luminance channels ^45,46^, which is thought to be primarily responsible for processing of pattern information ^18,19^. We then used the Image Calibration and Analysis Toolbox ^47^ in ImageJ to conduct granularity-based pattern analysis. In this process, widely used to study animal patterning ^48–51^, images are Fast Fourier band pass filtered into a number of granularity bands that correspond to different spatial frequencies. For each filtered image, the 舠energy舡 at that scale is quantified as the standard deviation of filtered pixel values and corresponds to the contribution to overall appearance from pattern elements of that size. Pattern energy spectra were calculated for each species in a comparison across 17 bandwidths (from 2 pixels to 512 pixels, by multiples of √2), which we used for both pairwise pattern comparisons (pattern maps can be created to visualize differences; Fig. 3a), and to categorize overall plumage pattern with PCA (Fig. 3c,d). Pattern difference values were calculated by summing absolute differences between energy spectra at each bandwidth ^47^, and principal components analyses revealed that the first three PCs explained ~93% of the variance in overall pattern energy (Fig. 3c; Extended Data Table 2) across species (Fig. 3d). Pattern PC1 has large positive loadings for most element sizes/granularities, indicating that species with high PC1 scores have numerous pattern elements of different sizes (whereas species with low PC1 are relatively homogenous and with little overall patterning; Fig. 3d). Pattern PC2 has large positive loadings for small pattern elements, and negative loadings for intermediate and large pattern sizes such that species with high PC2 scores have lots of small pattern components, and species with low PC2 scores have more intermediate and large pattern size contributions (Fig. 3d).

### Photographic quantification of woodpecker plumage colour and pattern

To validate the use of colour plates for quantifying meaningful interspecific variation in plumage colour and pattern among woodpeckers, we employed digital photographic and visual ecology methods to quantify the appearance of museum specimens and compared these results to those obtained using the colour plates. Specifically, we used ultraviolet and visible spectrum images to create standardized multispectral image stacks and then converted these multispectral image stacks into woodpecker visual space. Photos were taken with a Canon 7D camera with full-spectrum quartz conversion fitted with a Novoflex Noflexar 35 mm lens, and two Baader (Mammendorf, Germany) lens filters (one transmitting only UV light, one transmitting only visible light). We took profile-view photograph pairs (one visible, one UV) under full-spectrum light (eyeColor arc lamps, Iwasaki: Tokyo, Japan, with UV-coating removed), then converted these image stacks into woodpecker visual space using data from *Dendrocopos major* ^52^ and average visual sensitivities for other violet-sensitive bird species ^53^. The inferred peak-sensitivity (λmax) for the short-wavelength sensitive 1 (SWS1) cone of Great Spotted Woodpeckers, based on opsin sequence, is 405 nm ^52^. After generating images corresponding to the quantum catch values (i.e., stimulation of the different photoreceptor types), we performed granularity-based pattern analyses with the Image Calibration and Analysis Toolbox ^47^ in ImageJ ^41^ using the image corresponding to the stimulation of the avian double-cone, responsible for luminance detection ^45,46^, because this photoreceptor type is assumed to be involved in processing pattern information from visual scenes ^18,19^. Additionally, because relative stimulation values do not generate perceptually-uniform colour spaces ^54,55^, we implemented visual models ^56^ to generate Cartesian coordinates for the colour values from each of 1,000 randomly selected, 9-pixel diameter circles for each specimen and viewpoint (as we did with colour plates). Cartesian coordinates in this perceptually-uniform woodpecker colour space were then used to calculate pair-wise Mahalanobis distances ^44^ for each dyadic combination of measured specimens.

As with our colour plate-based analysis, we Z-score transformed colour and pattern distances (mean = 0, *SD* = 1), then combined these distances to create a composite plumage dissimilarity matrix incorporating overall plumage colour and pattern. Based on specimens available at the Cornell University Museum of Vertebrates, we endeavoured to measure up to three male specimens from at least one species of every woodpecker genus. We were able to measure 56 individuals from 23 woodpecker species (Extended Data Table 3). To compare the museum-based results to those from the colour plates, we derived species-level pairwise distances. We did so by finding the mean plumage distance between all specimens of one species and all of those of another, and repeating for all possible species pair comparisons. We repeated this process for both the colour only dissimilarity matrix, and the combined colour and pattern matrix. We subset the larger, plate-based colour-only and colour-plus-pattern matrices to the corresponding species, and compared the relevant matrices with Mantel tests. Our results from the museum specimens substantiated those from the illustrations舒we found tight correlations between colour dissimilarity (measured from specimens vs. measured from illustrations; *r* = 0.74, *p* < 0.001) and overall plumage dissimilarity (measured from specimens vs. measured from illustrations *r* = 0.72, *p* < 0.001).

### eBird data management, curation, and analysis

On 24 November, 2017 we queried the eBird database for all records of each of the 230 species in our final woodpecker phylogeny. We excluded records for which we had low confidence in the associated locality information. Specifically, we excluded: 1) historical records, which are prone to imprecise locality information and are not associated with effort information, 2) records from (0°, 0°), 3) records that were considered invalid after review by a human (thus, flagged but unreviewed records were included), and 4) records that came from transects of longer than 5 km in length. Because eBird has grown exponentially in recent years, we connected directly to the database to ensure maximal data coverage for infrequently reported species. We made this analytical decision because the automatic filters that flag unusual observations can be imprecise in regions of the globe infrequently visited by eBirders; flagged observations remain unconfirmed (and not included in products such as the eBird basic dataset) until they are reviewed, and backlogs of unreviewed observations exist in some infrequently birded regions. This approach allowed us to increase our sample size for infrequently observed species. In contrast, other species are very well represented in the database. To reduce downstream computational loads, we used the R package *ebirdr* (https://github.com/eliotmiller/ebirdr) to downsample overrepresented species in a spatially stratified manner. Specifically, for each of the 230 species, we laid a grid of 100 x 100 cells over the species’ extent, and randomly sampled and retained 60 points per cell. For most species, this had little to no effect, and fewer than 10% of points were thinned and removed from analysis; for a small number of well-sampled North American species, this excluded over 90% of points from analysis (Extended Data Table 4). In sum, this process reduced the original dataset from 13,513,441 to 1,037,628 records.

We used the R package *hypervolume* ^57^ to create pseudo-range maps around each species’ point locations. Hypervolumes account for the density in the underlying points and can have holes in them, and are therefore much better suited to describing species’ ranges than are, e.g., minimum convex polygons ^57^ For every dyadic comparison (i.e., for every species pair comparison), we used *hypervolume* to calculate the Sørenson similarity index between the species’ inferred geographic ranges. We summarized these similarities in a pairwise matrix, which we subsequently converted to a dissimilarity matrix such that a value of 1 represented complete allopatry (no overlap in geographic distributions), and a value of 0 represented perfect sympatry (complete overlap in geographic distributions).

We used the *raster* package to match each species’ point locations to climatic data at each point using WorldClim bioclimatic data ^58^. These data describe the annual and seasonal climatic conditions around the globe. After querying species’ climatic data, we bound the resulting files together and ran a single large correlation matrix PCA across all climate variables except bio7, which is simply the difference between bio5 and bio6. We retained species’ scores along the different PC axes, and used scores along the first two PC axes to calculate species-level hypervolumes in climate space. These first two axes explained 85% of the variance in the climates occupied by woodpeckers. The first axis described a gradient from places that are warm in general, and that are not cold in winter, to areas that show seasonal variation in temperature and large diurnal shifts in temperature. The second axis described a gradient from areas that receive precipitation in seasonal pulses, have some hot months and have large swings in temperature over the course of a day, to areas that always receive lots of rain. Again, for each dyadic comparison, we calculated a Sørenson similarity index, and then converted the resulting values to a dissimilarity matrix.

### Querying habitat data

We used *ebirdr,* which harnesses GDAL (http://www.gdal.org/), to bind species’ point locations into ~50 MB-sized tables, then converted the resulting tables into KML (Keyhole Markup Language) files, which we uploaded and converted into Google Fusion Tables (https://fusiontables.google.com). This particular file size was chosen after we employed a trial- and-error process to determine the most efficient query size for Google Earth Engine (see below). Once accessible as a Fusion Table, we fed the tables into custom Google Earth Engine scripts. For every eBird observation, these scripts identified the MODIS satellite reflectance values ^59^ from the observation location within a 16-day window of the observation. We queried data specifically from the MODIS MCD43A4 Version 6 Nadir Bidirectional reflectance distribution function Adjusted Reflectance (NBAR) data set, a daily 16-day product which 舠provides the 500 meter reflectance data of the MODIS 舖land’ bands 1-7 adjusted using the bidirectional reflectance distribution function to model the values as if they were collected from a nadir view舡 (https://lpdaac.usgs.gov/node/891). At the time of query, this dataset was available for the time period 18 February 2000 to 14 March 2017, which corresponded to the time period in which most of our eBird records were recorded. The year of all other records was adjusted up or down to fall within the available satellite data, e.g., observations from 10 November 2017 became 10 November 2016. This method is appealing in that it incorporates species’ spatiotemporal variation in habitat availability and use, although for most woodpecker species such variation is minimal.

After querying species’ habitat data, we downloaded and combined the resulting files from Google Earth Engine, dropping any records that were matched to incomplete MODIS data. *ebirdr* contains functions to automatically combine and process these files from Google Earth Engine. We then ran a single large correlation matrix PCA across all 7 MODIS bands. Before doing so, we natural log-transformed bands 1, 3, and 4, as a few extreme values along these bands hampered our initial efforts to ordinate this dataset. We retained the first two PC axes, which explained 81% of the variance in the habitats occupied by woodpeckers. The first described a gradient from closed forests to open, reflective habitats. The second described a gradient from regions with high visible and low infrared reflectance to those with low visible and high infrared reflectance. This dichotomy is used to identify snow in MODIS snow products (https://modis.gsfc.nasa.gov/data/atbd/atbd_mod10.pdf). Thus, at the species-average level, the second habitat PC axis functionally described a gradient between seasonally snow-covered (temperate) forests and tropical woodland. Again, for each dyadic comparison, we calculated a Sørenson similarity index, and then converted the resulting matrix to a dissimilarity matrix.

### Multiple distance matrix regression

After the steps described above, we had data of four variables hypothesized to explain plumage variation across woodpeckers in the form of four pairwise distance matrices: genetic distances, climate dissimilarity, habitat dissimilarity, and geographic range dissimilarity. We combined the plumage colour and the plumage pattern dissimilarity matrices into a single matrix by independently standardizing each using z-scores, then calculating the element-wise sum of each dyadic comparison. We then related the four explanatory matrices to the single dependent plumage dissimilarity matrix using multiple distance matrix regression, with 999 permutations ^60^. The resulting model was highly significant (*p* = 0.001), but fairly low in explanatory power (*r*^2^ = 0.03), reflecting the massive variation incorporated into these five 230 x 230 matrices. Three of the four dissimilarity matrices were significantly and positively associated with plumage dissimilarity: increasing genetic distance (*p* = 0.02), climate dissimilarity (*p* = 0.007), and habitat dissimilarity (*p* = 0.006) all lead to increasing plumage dissimilarity. At this scale, geographic range dissimilarity was not significantly associated with plumage dissimilarity.

### Modified Mantel correlogram

The likelihood that sympatric species evolve plumage mimicry is not thought to be monotonically related to range overlap. Instead, hypotheses to explain plumage mimicry propose that only certain species pairs that are both closely sympatric and ecologically similar will converge dramatically in appearance ^14^ Thus, we did not expect a continuous relationship between geographic range dissimilarity and plumage dissimilarity; rather, we expect plumage convergence in only dyads with high geographic overlap (greater than some threshold). We therefore implemented a modified Mantel correlogram approach to test whether such a threshold existed ^61^. For this, we manually created a series of matrices where we converted all elements in the plumage dissimilarity matrix to values of 1, except for dyads with plumage dissimilarity scores in a certain range. Specifically, the dissimilarity scores within a given range for a given analysis (ranges tested: 0-0.1, 0.1-0.2, 0.2-0.25, 0.25-0.3, 0.3-0.35, 0.35 0. 4, 0.4-0.5, 0.5-0.6, 0.6-0.7, 0.7-0.8, 0.8-0.9, and 0.9-1) were set to a value of 0, and all other dissimilarity scores remained unmanipulated. We then sequentially input these matrices as the dependent variable into the same multiple distance matrix regression described above, repeatedly calculating the significance and partial correlation coefficient of the geographic range dissimilarity matrix with that of plumage dissimilarity. This approached allowed us to examine how the correlation between plumage and range dissimilarities varied across a range of plumage dissimilarities, while simultaneously incorporating the influences of evolutionary relationships, and climate and habitat dissimilarities.

We found that geographic range dissimilarity was significantly associated with plumage dissimilarity for the most similar looking species pairs (Extended Data Fig. 1). Dyadic comparisons with plumage dissimilarities of 0-0.2 include such pairs as *Picoides pubescens* and *Picoides villosus* (purported plumage mimics), *Gecinulus grantia* and *Blythipicus pyrrhotis (which are quite similar looking*), and *Picus awokera* and *Melanerpes striatus* (not closely similar, but do share colour and pattern elements). Notably, the relationship was reversed at intermediate levels of plumage dissimilarity; species pairs that phenotypically differed by 25-35% were associated with partial but incomplete range overlap. Dyadic comparisons with dissimilarities in this range include *Campethera abingoni* and *C*. *maculosa* and *Veniliornis spilogaster* and *Celeus obrieni.* This pattern mirrors recent results showing the greatest degree of plumage differentiation between pairs of birds at intermediate levels of sympatry ^62^. The fact that some degree of sympatry is associated with rapid plumage divergence is expected by theory ^63^, and is likely due to strong selection to avoid unsuccessful hybridization (i.e., reinforcement), or to avoid accidentally targeting heterospecifics for aggression ^64^. Whether the relaxation of plumage divergence in closer sympatry could be attributed to shared habitats or climates, or to some other selective pressure, was heretofore unknown ^62^. We show that, remarkably, in woodpeckers, after accounting for other likely selective forces, either one or both of the species in pairs that have attained close sympatry may evolve towards the phenotype of the species with which they co-occur.

### Identification of putative plumage mimics

We developed a novel and powerful method to identify high-leverage dyadic comparisons in Mantel tests and multiple distance matrix regressions. We used this to identify species pairs that have converged above and beyond that expected by shared climates and habitats. The process works as follows. In the first step, the observed correlation statistic is calculated. In our case, that was the correlation coefficient of a thresholded plumage dissimilarity matrix (values from 0-0.2 set to 0, all others set to 1) with the geographic range dissimilarity matrix. The statistic can also be the correlation coefficient from a regular or partial Mantel test; we confirmed that the method yielded similar results when we employed it with a partial Mantel test between the continuous plumage dissimilarity matrix, geographic range dissimilarity, and genetic distance. In the second step, each element (dyad) in the relevant matrix is modified in turn, and the relevant correlation statistic calculated and retained after each modification. We tested three methods of modifying dyads, i.e., three different approaches to this second step. All yielded similar results. (A) The value can be randomly sampled from the off-diagonal elements in the matrix. (B) The value can be set to NA and the correlation statistic calculated using all complete observations. (C) For the thresholded matrix, the test element can be swapped for the other value; zeros become ones, and ones become zeros. In the third step, only necessary for approach A, the process is iterated multiple times, and the modified correlation coefficient for every element, at each iteration, is stored as a list of matrices. In the fourth step, again only necessary for approach A, these matrices are summarized by taking the element-wise average. In the fifth step, the leverage of each dyad is calculated by subtracting the observed correlation statistic from each element in the averaged matrix. Finally, the matrix can be decomposed into a pairwise table and sorted by the leverage of each dyad. In our case, dyads that have high leverage, and are large contributors to the positive correlation between close plumage similarity and geographic range overlap, have the largest negative values (i.e., modifying their observed plumage dissimilarity score most diminished the observed positive correlation between range and plumage).

We used this method to identify the most notable plumage mimics across woodpeckers, after accounting for shared evolutionary history, climate, and habitat use. Many purported mimicry complexes were responsible, including the Downy-Hairy system ^21^, and repeated convergences between members of *Veniliornis* and *Piculus* ^22^, *Dinopium* and *Chrysocolaptes*, and *Dryocopus* and *Campephilus* ^23^. Convergence between the Helmeted Woodpecker (*Dryocopus=Celeus galeatus*) and *Campephilus robustus* was also detected ^15^, as was convergence between members of *Thripias* and *Campethera*, *Meiglyptes* and *Blythipicus,* and *Hemicircus* and *Meiglyptes.*

### Phylogenetic least squares regression

We derived species’ average scores along the first two axes of a plumage colour principal components analysis (PCA, Fig. 2), a plumage pattern PCA (Fig. 3), the climate PCA described above, the habitat PCA described above, and species’ average latitude (absolute value) and longitude of distribution. Additionally, we mined body mass data from Dunning (2007). For those species for which mass was listed separately for males and females, we calculated sexual size dimorphism *sensu* Miles et al. (2018). These authors additionally reported dimorphism measures from a number of species not available in Dunning (2007). We then combined these datasets, resulting in sexual size dimorphism measures for 94 of 230 species. During this process, we noticed that one of the most well-known of sexually size-dimorphic species, *Melanerpes striatus*, was characterized in both databases as having larger females than males. This is incorrect舒males are much larger than females舒and we replaced the values with the midpoint of ranges given in del Hoyo et al. (2017). We used *Rphylopars* ^67^ to impute missing body mass and size dimorphism data, which we did using a Brownian motion model and the observed variance-covariance matrix between all traits except for plumage colour and pattern.

Treating climate, habitat, latitude, longitude, natural log body mass, and sexual size dimorphism as explanatory variables, we used multi-model inference to identify phylogenetic generalized least squares (PGLS) regression models that explained each of the four PCA plumage axes of interest. We also visualized pairwise correlations and distributions of these traits using *corrplotter* ^68^ (Extended Data Fig. 2). We used a model averaging approach to determine which explanatory variables strongly influenced plumage (Fig. 4).

### Code availability

All computer code necessary to run these analyses is available in the purpose-built R package *ebirdr,* available at https://github.com/eliotmiller/ebirdr.

## Acknowledgements

We thank T. Auer, D. Caetano, D. Fink, R. Grant, L. Harmon, W. Hochachka, K. Horton, S. Kelling, R. Maia, A. Patton, M. Pennell, M. Strimas-Mackey, D. Toews, J. Uyeda, C. Wood, and R. Zenil-Ferguson for statistical, graphical, and intellectual input. ETM, GML, and RAL were supported by NSF grants 1402506, 1523748, and 1523895, respectively. ETM, GML, RAL, and ACL were supported by the Cornell Lab of Ornithology. BGF was supported by Banting Canada (379958) and the Biodiversity Research Centre at the University of British Columbia.

## Author contributions

ETM, GML, BGF, ACL, and RAL were responsible for conceptualization and writing of the manuscript. ETM and RAL were responsible for analysis and visualization.

## Data deposition statement

All data is available in the manuscript or the supplementary materials. Reprints and permissions information is available at www.nature.com/reprints

## Competing interests

The authors declare no competing interests.

Correspondence and requests for materials should be addressed to

---

**Extended Data Table 1.** Principal components analysis of 230,000 CIELAB color values (1,000 samples from each of 230 woodpecker species). The first two PCs explain ~82% of the variance in color. PC1 has high loading values for luminance (L*) and yellowness (positive values of b*), and PC2 has high loading values for greenness (negative values of a*).

|  | PC1 | PC2 | PC3 |
| --- | --- | --- | --- |
| *Eigenvalues* | 1.17 | 1.04 | 0.74 |
| *Proportion of Variance* | 0.46 | 0.36 | 0.18 |
| *Cumulative Proportion of Variance* | 0.46 | 0.82 | 1 |
| Color variable | *PC loadings* |  |  |
| L* | **0.70** | 0.29 | -0.65 |
| a* | 0.01 | **-0.92** | -0.39 |
| b* | **0.71** | -0.27 | 0.65 |

**Extended Data Table 2.** Principal components analysis of pattern energy spectra across 230 species of woodpeckers. The three PC axes explain ~93% of the variance in pattern across all species.

|  | PC1 | PC2 | PC3 |
| --- | --- | --- | --- |
| Standard deviation | 3.08 | 2.12 | 1.34 |
| Proportion of Variance | 0.56 | 0.26 | 0.11 |
| Cumulative Proportion | 0.56 | 0.82 | 0.93 |

**Extended Data Table 3.**
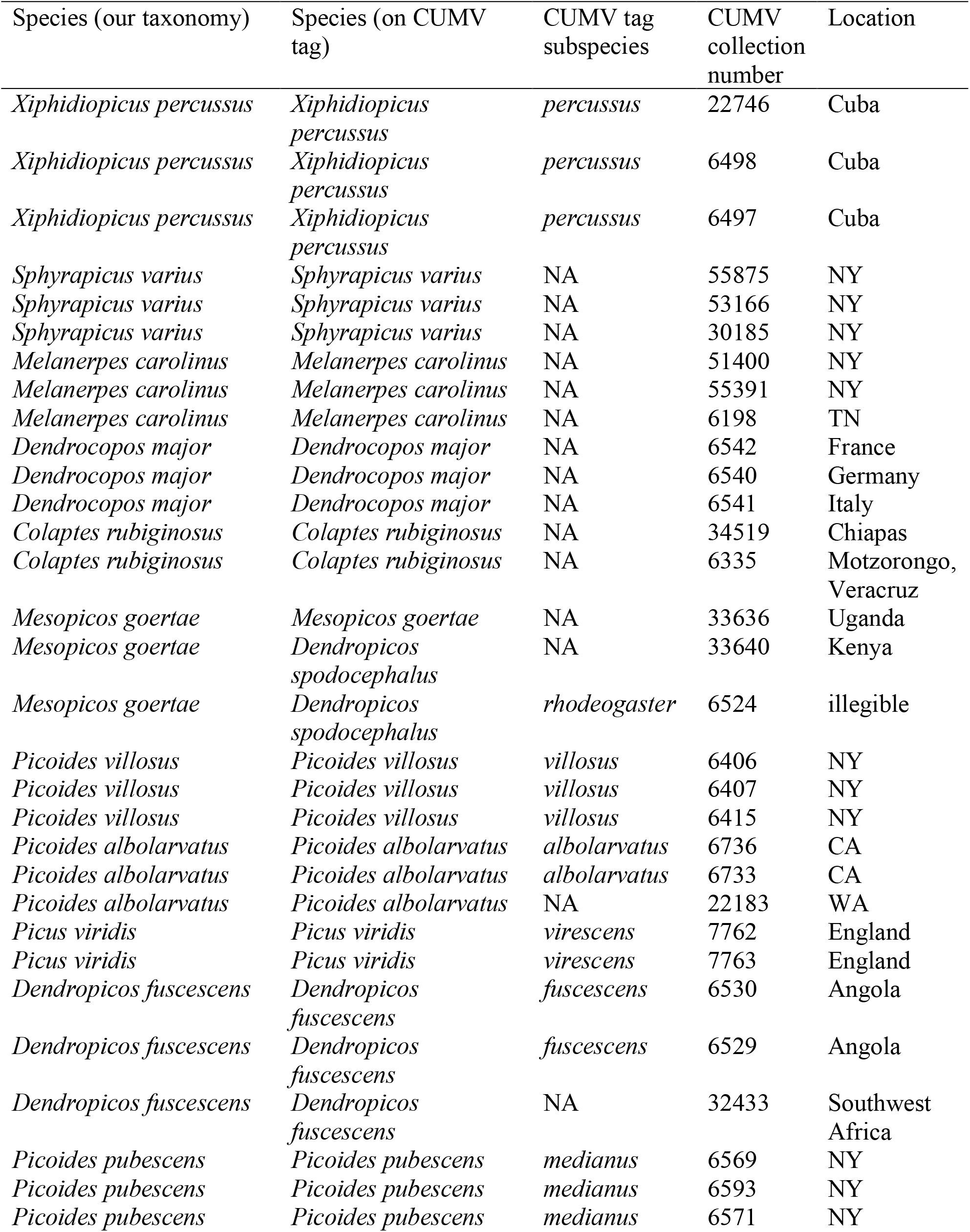
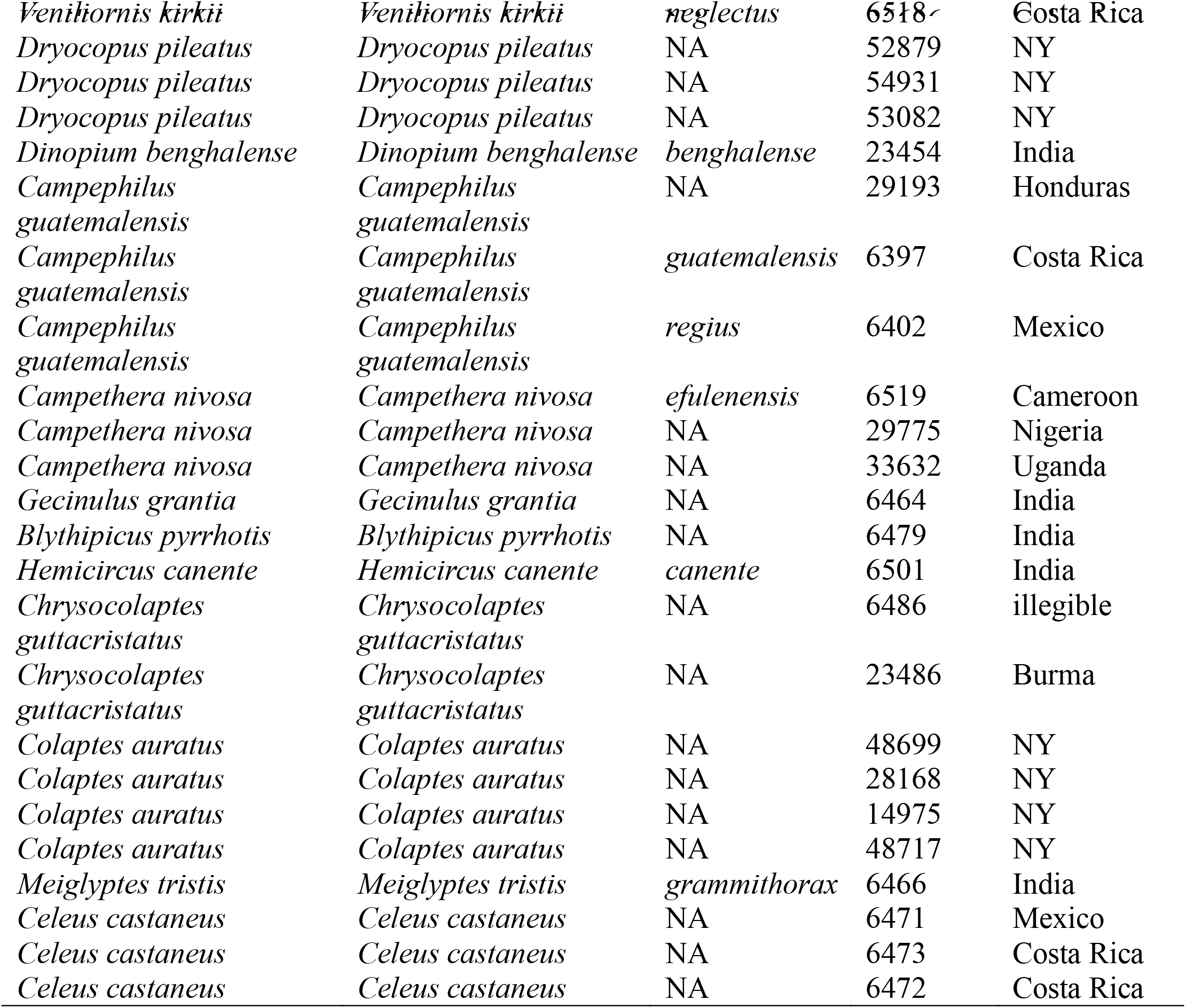
Synopsis of specimens from the Cornell University Museum of Vertebrates (CUMV) used for color and pattern analyses.

**Extended Data Table 4.**
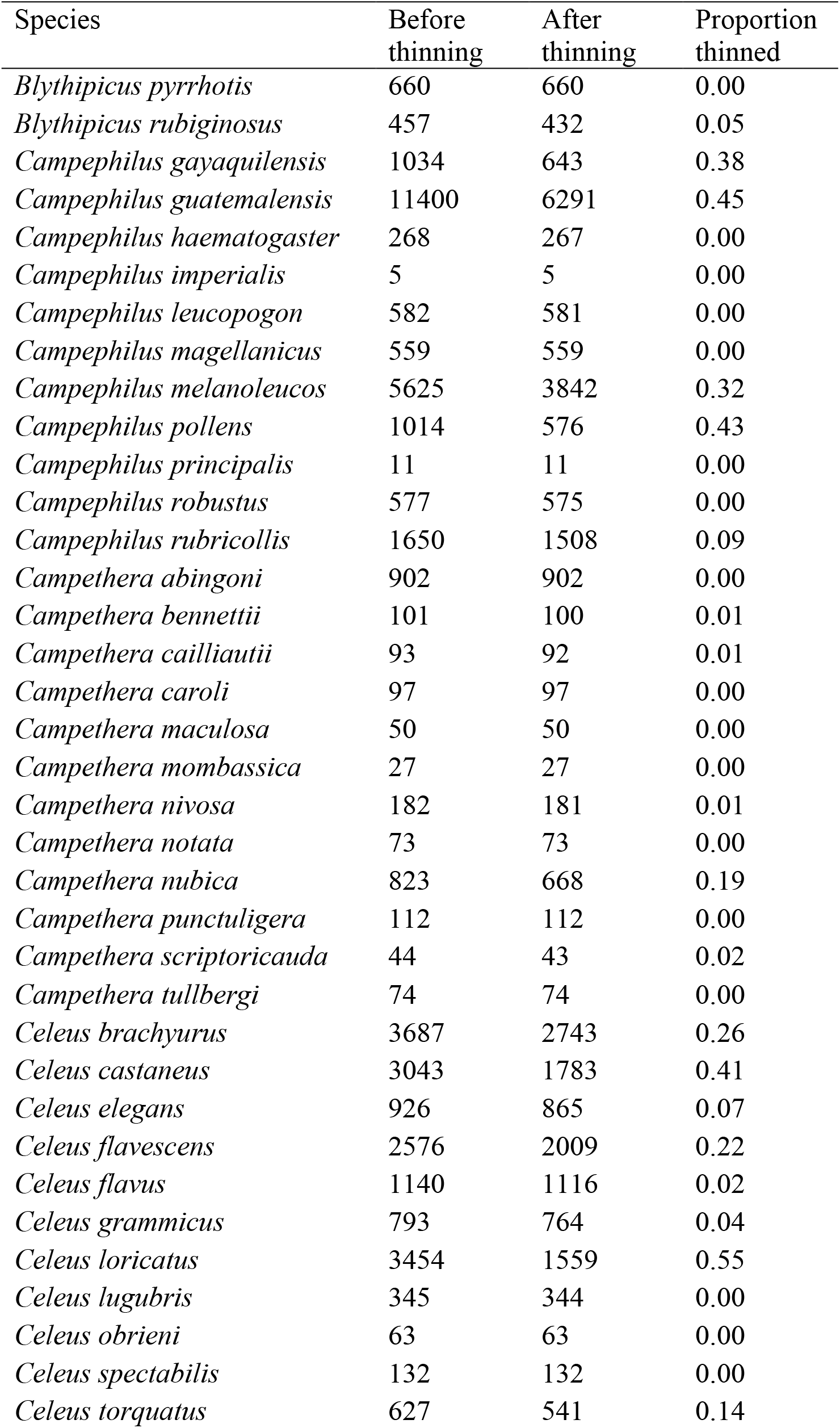
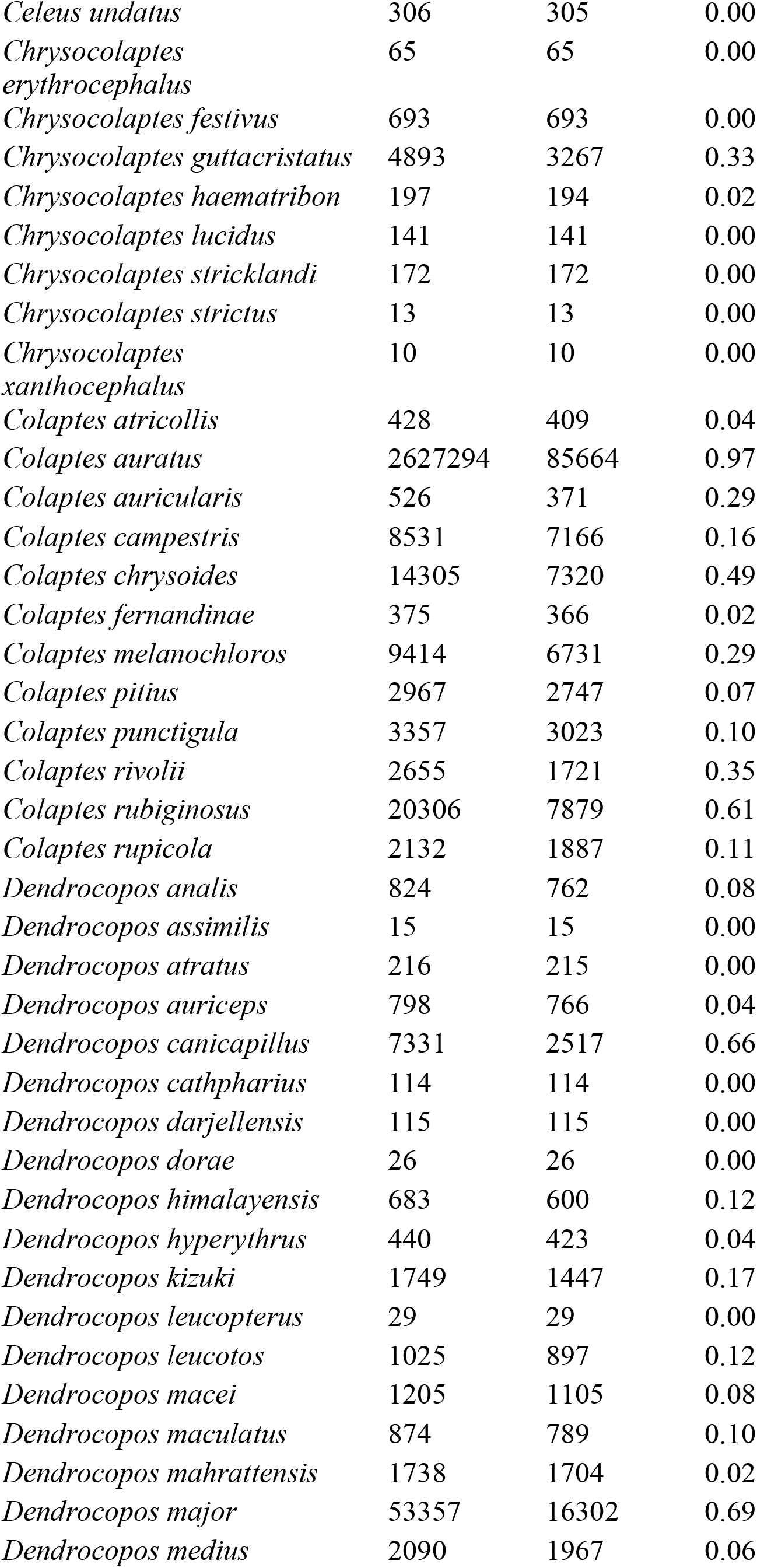
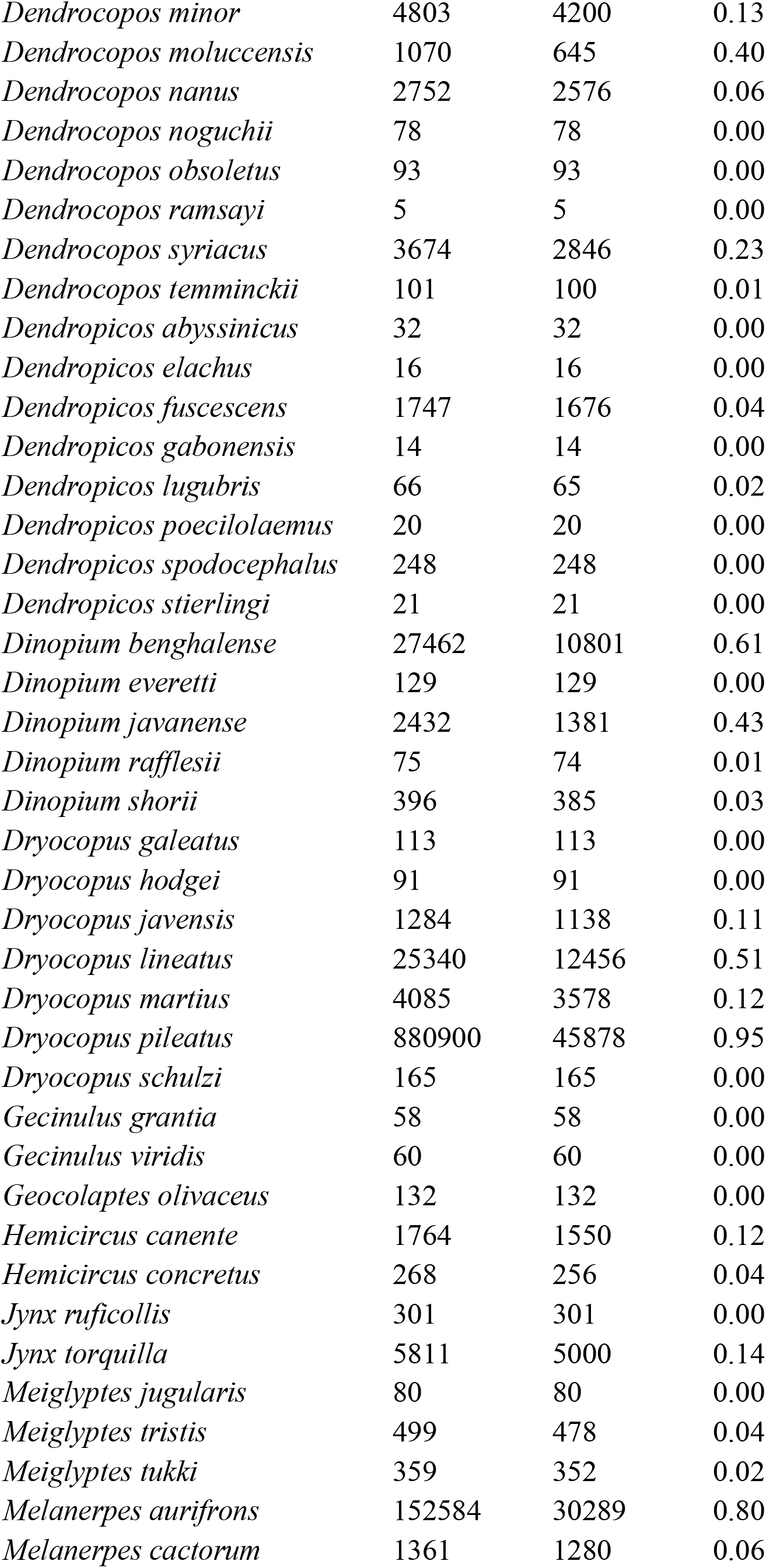
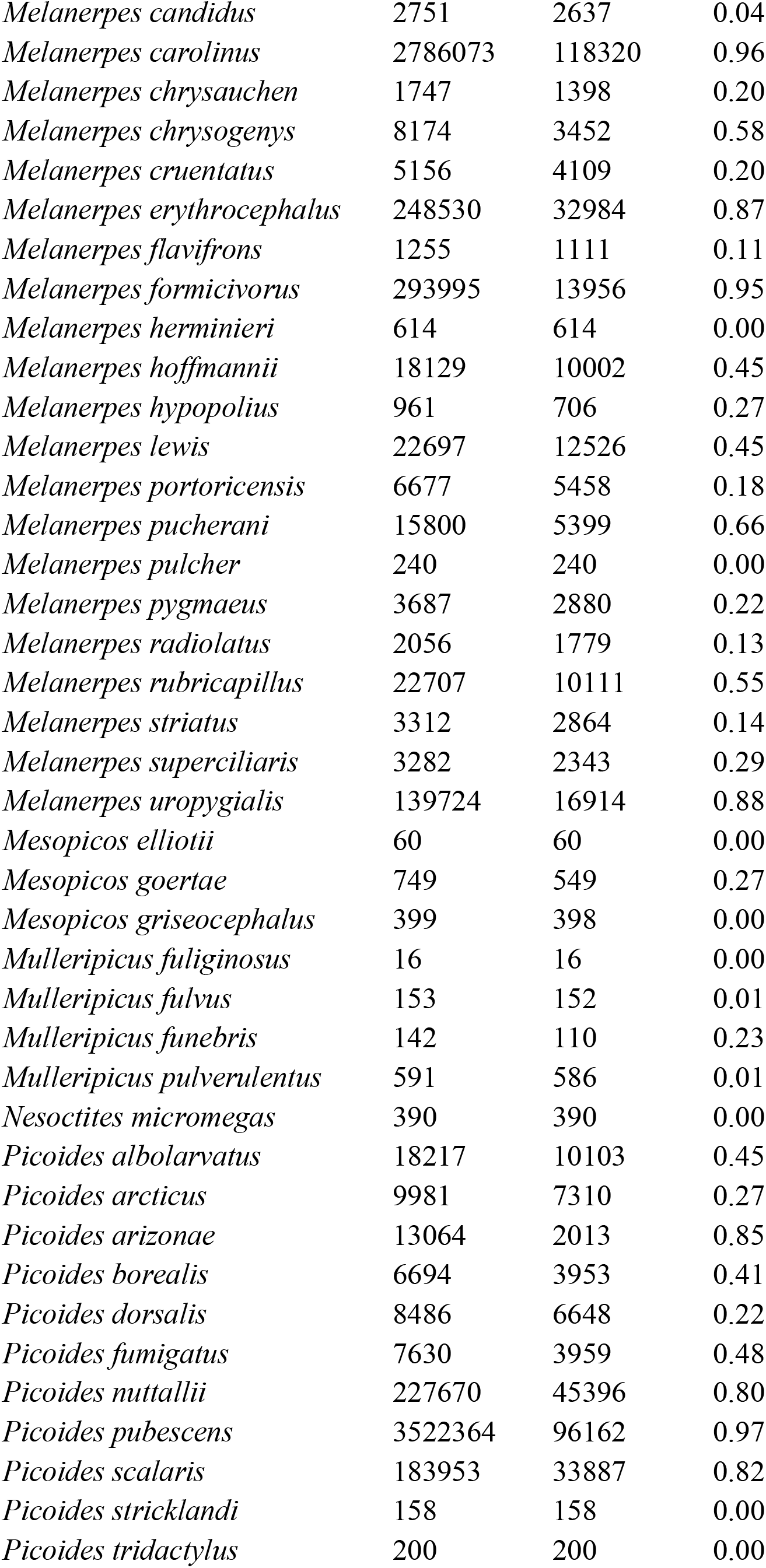
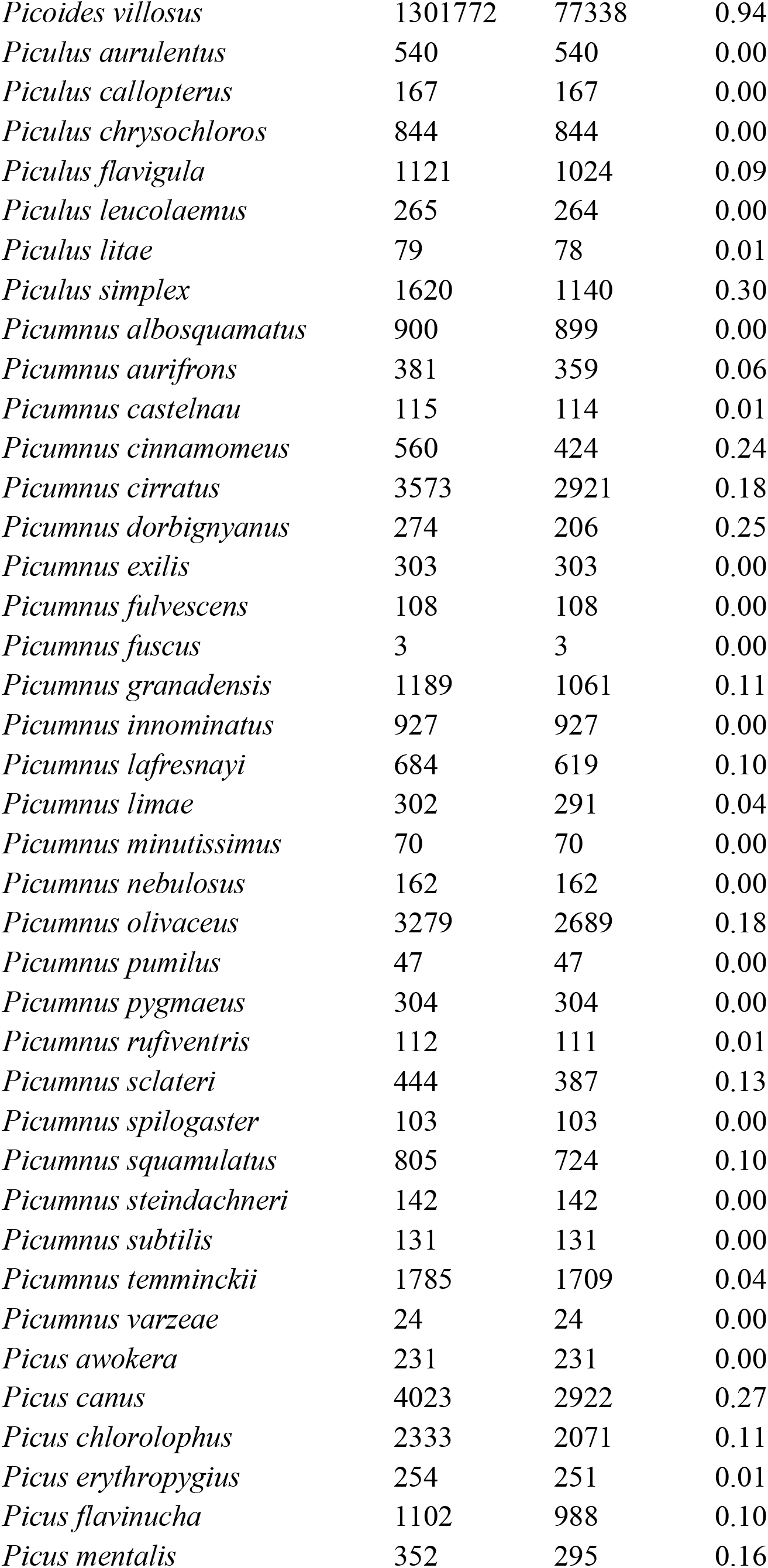
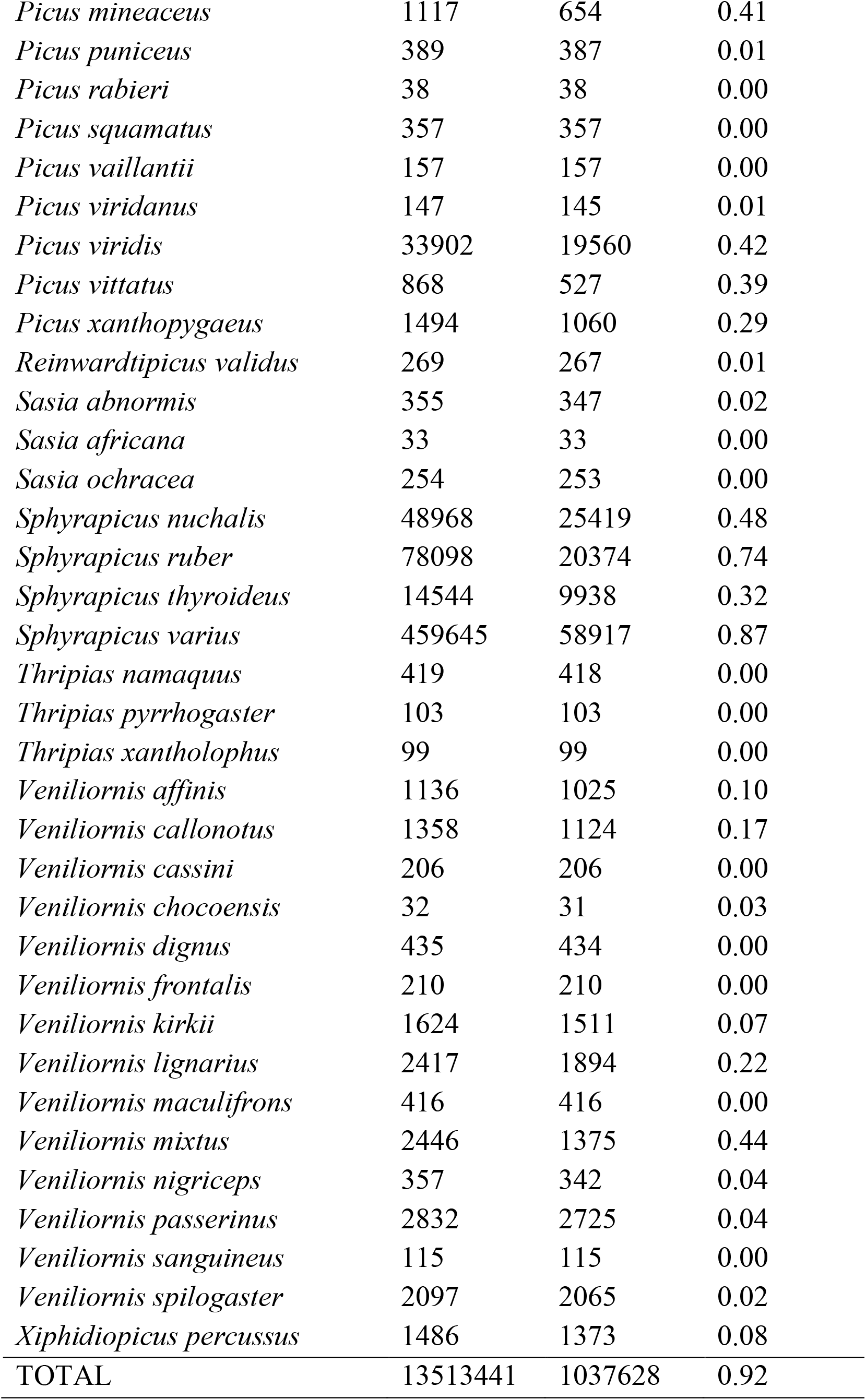
Species-level sample sizes of eBird point locations before and after spatial downsampling.

**Extended Data Fig. 1.**
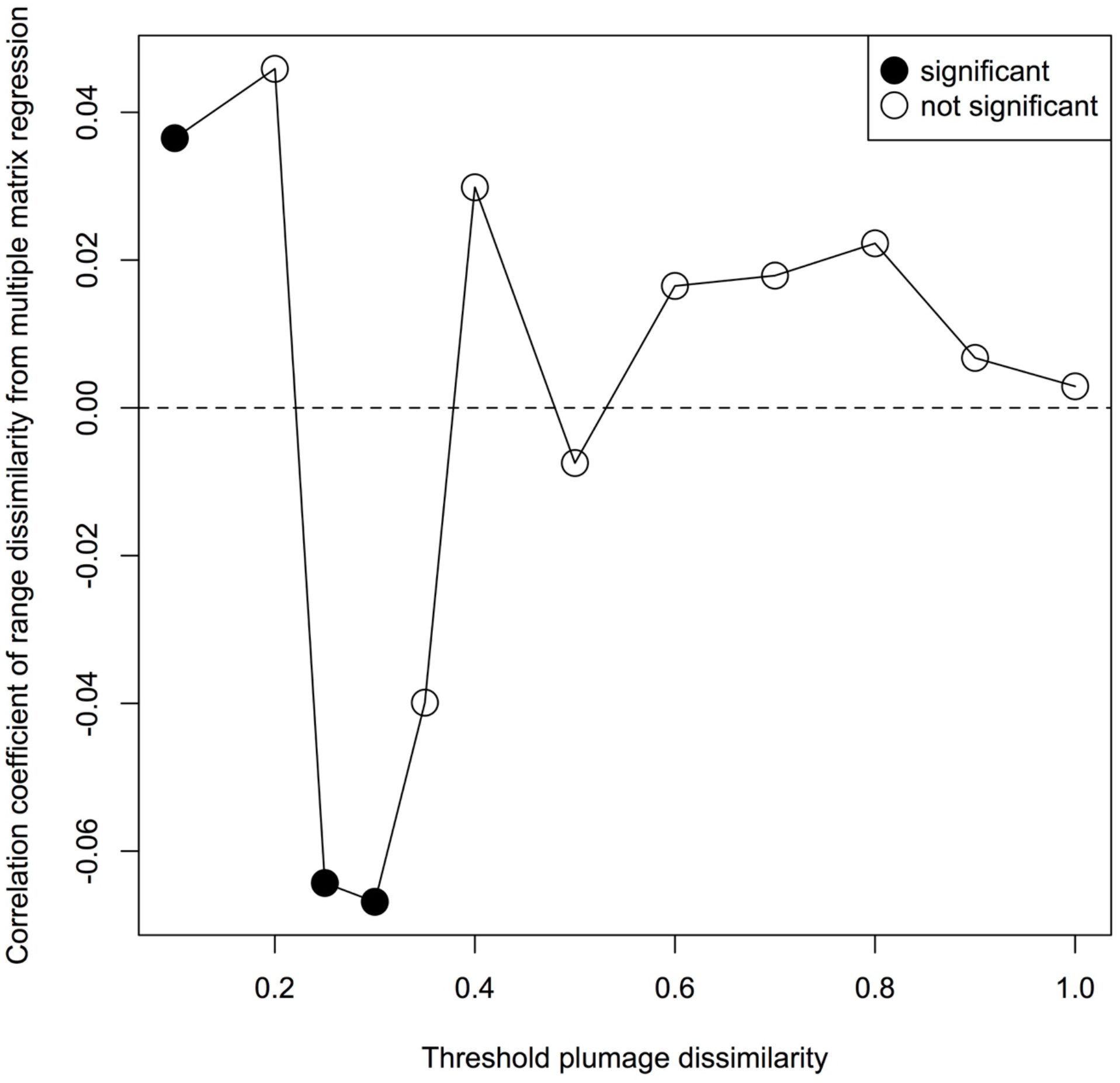
Illustration of the results of the modified Mantel correlogram approach. Plumage dissimilarity is positively associated with range dissimilarity for the most similar-looking, geographically overlapping species pairs. Additionally, this illustration of results from a modified Mantel correlogram approach reveals that the two variables are negatively associated with one another for species dyads of intermediate plumage similarity.

**Extended Data Fig. 2.**
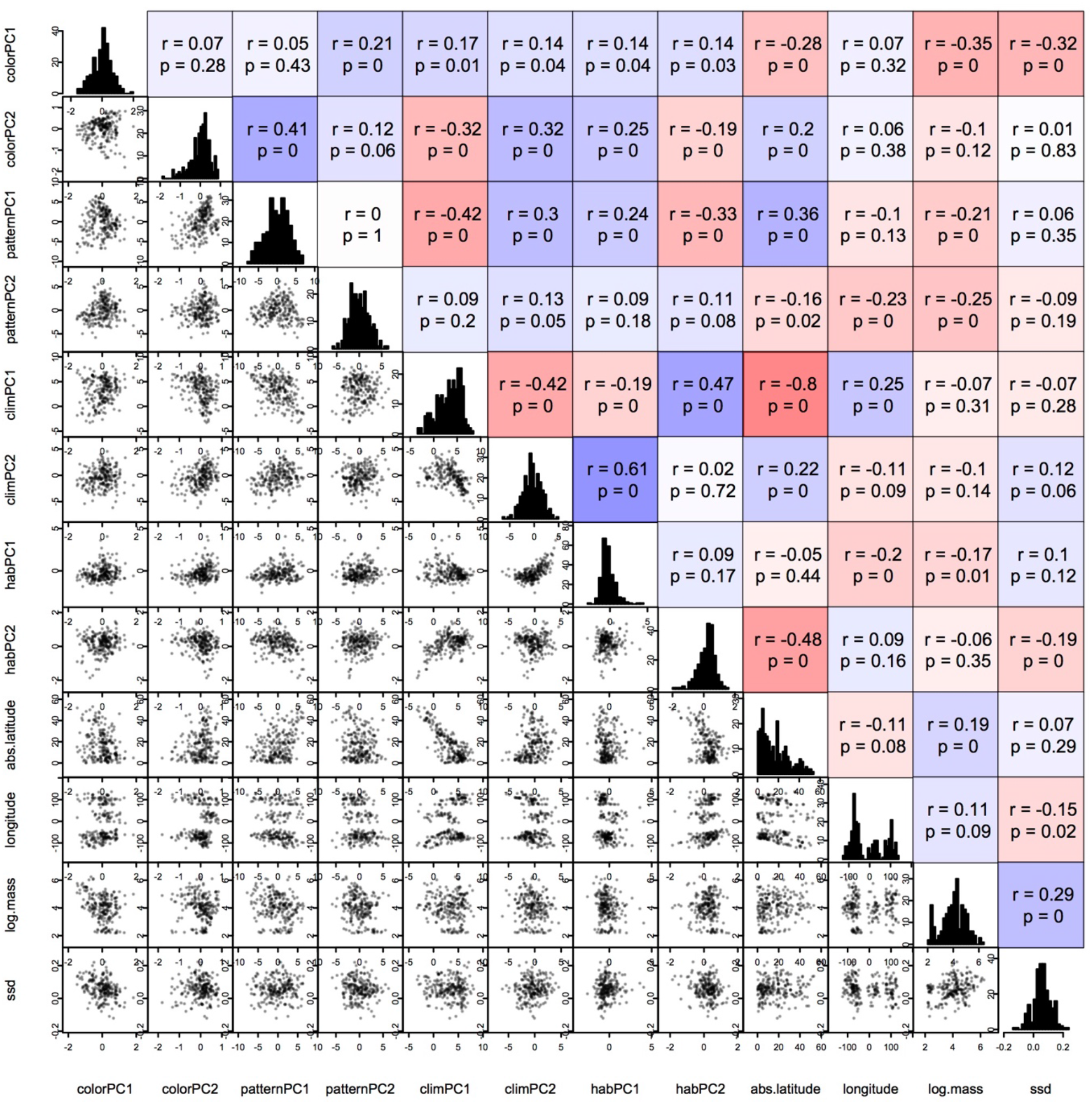
Paneled correlation plot showing the relationships between all species-level averaged traits. Climate, habitat, latitude, and woodpecker body mass and plumage appearance exhibit a variety of intercorrelations. The lower triangle panels show the pairwise trait relationships. The diagonal panels contain histograms of the indicated variable. The upper triangle panels summarize the Pearson’s correlation coefficient and the significance of the relationship, both numerically and in color, where bright red indicates a strong negative correlation, and bright blue indicates a strong positive correlation.

